# One thousand samples per day capillary-flow LC/MS/MS for high-speed, high-sensitivity and in-depth proteomics

**DOI:** 10.1101/2023.06.05.543682

**Authors:** Ayana Tomioka, Ryota Tomioka, Issei Mori, Yosuke Isobe, Makoto Arita, Koshi Imami, Eisuke Kanao, Kosuke Ogata, Yasushi Ishihama

## Abstract

We developed a capillary-flow LC/MS/MS system with ultrahigh speed, enabling a throughput of 1,000 samples per day while maintaining high sensitivity and depth of analysis. In targeted LC/MS mode, 36 endogenous phosphopeptides in HeLa cells, including EphA2- derived phosphopeptide isomers, were successfully quantified with high selectivity and linearity by combining ion mobility separation. When 500 ng of HeLa cell digest was measured 100 times repeatedly in data-dependent acquisition mode, the coefficient of variation of retention time, peak intensity and number of identified peptides were on average 3.4%, 19.8%, and 6.0%, respectively. In data-independent acquisition mode, this system achieved the identification and quantification of 3,139 protein groups from a 100 ng HeLa cell digest and 2,145 protein groups from a sample of only 10 ng. The coefficient of variation of protein commonly quantified in the triplicate analysis ranged from 12 to 24% for HeLa digest samples ranging from 10 to 1000 ng. Finally, we applied this high-speed system to the spatial proteomics of the mouse brain, and succeeded in capturing the proteome distribution along a 96-sectioned brain structure in 135 minutes. This is the first LC/MS/MS system to achieve both more than 500 samples per day and more than 3000 identified protein groups ID with less than 100 ng human cultured cells simultaneously.

## Introduction

Proteins play a crucial role in biological processes inside and outside cells by strictly regulating various biological functions^1^. Despite their significance, many of these intricate biological mechanisms remain largely unknown. One promising approach for unraveling these mysteries is shotgun proteomics, which enables comprehensive analysis of protein profiles in living organisms^2^. With the advancement of shotgun proteomics, it is now possible to simultaneously analyze over 10,000 human proteins in a single measurement ^3, 4^, providing comprehensive insights not only into changes in protein expression but also into various proteoforms^5^ due to genetic variations, alternatively spliced RNA transcripts and post-translational modifications^6, 7^.

The need for higher sensitivity and faster liquid chromatography/tandem mass spectrometry (LC/MS/MS) in shotgun proteomics is increasing in a variety of applications. For example, in biomarker discovery using clinical samples, the low throughput of currently used nano-flow LC/MS/MS limits the total number of samples that can be analyzed to a few hundred to a few thousand^8^. In addition, spatial proteomics, which aims to simultaneously obtain spatial information on thousands of proteins from hundreds of micro-sections, requires weeks to a month to analyze hundreds or more tissue sections, making current methods impractical^9, 10^. Furthermore, even in single-cell proteome analysis, which has shown remarkable progress in the past few years^11, 12^, at least several hundred cells must be analyzed to discover heterogeneity in various cell types, and the throughput of current systems does not meet this need. Nevertheless, in single-cell analysis, improving the sensitivity of nano-flow LC/MS/MS should be a top priority ^11, 12^ and a recent study has shown that combining a gradient elution method at an ultra-low flow rate of 100 nL/min for 35 min with a high-sensitivity mass spectrometer can provide a high sensitivity of up to 2,083 identified proteins per single cell^13, 14^. However, as long as the current nano-flow LC/MS/MS system is used, it is not easy to achieve even higher throughput while maintaining high sensitivity. On the other hand, for large sample analyses, such as clinical samples, efforts are focused on improving the throughput of LC/MS/MS measurements^8, 15^. Several studies have demonstrated the robustness of micro-flow LC/MS/MS systems with flow rates ranging from 50 to 800 μL/min and 2.1 mm inner diameter columns^16–20^. One of these systems has successfully identified thousands of protein groups and has been applied to the classification of COVID-19 infections and consecutive measurements^20^. A report of a 50 μL/min flow rate, 30-min gradient analysis identified 3,210 protein groups in a 1 μg HeLa digest^17^. Another report using a short gradient time at 800 μL/min identified 1,937 protein groups in a 0.5 min gradient method and 5,004 protein groups in a 5 min gradient method for a 5 μg K562 digest^18^. In other reports, 5,286 or 5,844 proteins were identified from 3 μg K562 digest using a 3- or 5-minute gradient elution at a flow rate of 500 μL/min, respectively^19^. These reported micro-flow LC/MS/MS systems represent good successes in terms of system robustness and LC/MS/MS speed, enabling measurements as fast as 400 samples per day (SPD), but they require sample amounts on the microgram scale of 1 to 10 μg, due to a loss of sensitivity^8^.

In recent years, researchers have focused on capillary-flow LC with 1 - 10 μL/min to accelerate LC/MS/MS while maintaining sensitivity comparable to nano-flow LC with less than 1 μL/min ^21–24^. Capillary-flow LC reduces the time required for sample loading, column washing, and column equilibration compared to nano-flow LC. To address the challenge of faster LC/MS/MS, studies have successfully shortened the LC cycle time by increasing the flow rate only during sample loading^21, 23, 24^ and by utilizing dual-trap columns in well-designed LC configurations^22^. These approaches have achieved high sensitivity with sample amounts as low as 5 ng and have identified 1074 protein groups at a rate of 200 SPD^24^. However, while these methods have increased sensitivity, their throughput remains lower than that of micro-flow LC/MS/MS systems. An alternative way to achieve high-throughput measurements in direct-infusion mode has also been proposed^25^. This infusion mass spectrometry without LC, combined with the SILAC method, successfully measured 132 samples within 4.4 hours. While this method has excellent throughput, the complexity of the peptide mixture limits the depth of analysis, resulting in the identification of 525 proteins in 2.5 min of MS data collection.

As mentioned above, while various efforts have been made to increase the sensitivity and throughput of proteome measurement, it has been difficult to achieve both of these goals simultaneously. However, since the samples actually used in high-speed proteome measurements include single cells, small tissue fragments, and minute clinical specimens, it is essential to achieve both throughput and sensitivity.

In this study, we have developed an unprecedented high-throughput analysis, “machine-gun proteomics,” that can analyze up to 1,000 SPD without compromising sensitivity and depth of proteome analysis. By setting an upper limit for LC flow rate that does not sacrifice sensitivity based on the relationship between flow rate and electrospray ionization (ESI) efficiency, and combining it with a sub-minute gradient, we have achieved both high speed and sensitivity comparable to nano-flow LC/MS/MS. In addition, minimizing LC dead volume and incorporating trapped ion mobility spectrometry (TIMS)^26–28^ as an additional separation dimension expanded the separation efficiency and overcame the trade-off between sensitivity, throughput, and depth of analysis, enabling 1000 SPD with 3000 protein quantification from 100 ng HeLa digested cells.

## Results

### Relationship between ESI efficiency and flow rate

In shotgun proteomics, nano-flow LC/MS/MS utilizing nanoESI interface has been widely used to achieve high resolution in LC together with high sensitivity in MS. However, the use of nano-flow LC requires considerable time for sample loading and column washing and equilibration, resulting in longer LC cycle time between injections. Thus, there is a trade-off between measurement throughput and sensitivity, and it is necessary to know the exact upper limit of flow rate that is compatible with sensitivity in order to accelerate LC/MS/MS. Therefore, we investigated the extent to which ionization efficiency decreases as flow rate is increased from nano-flow (< 1 µL/min) to capillary-flow (1 - 14 µL/min) and to micro-flow (> 50 µL/min). Note that the flow range defined in this study is based on the optimum flow rate depending on the typical column diameters (nano-flow < 150 µm ID, capillary-flow < 0.53 mm ID and micro- flow > 1 mm ID).

To eliminate the concentration effect by the LC column and to evaluate only the relationship between flow rate and ionization efficiency, measurements were performed by direct infusion mass spectrometry. Three synthetic peptides derived from trypsin-digested albumin (Bovine) were used as model peptides and prepared so that the infused amount of peptide per time was equal (Supplementary Table S1). When the intensity at a flow rate of 250 nL/min, typical flow rate of nano-flow LC/MS/MS with 75 µm diameter column was set to 100%, the sensitivity decreased to 27% at 10 µL/min and to 2.3% at 200 µL/min (Fig. 1a and Supplementary Table S1). In other words, to obtain the same sensitivity as nano-flow LC/MS/MS, a sample amount of 4 times more at 10 µL/min and 50 times more at 200 µL/min is required.

**Figure 1.**
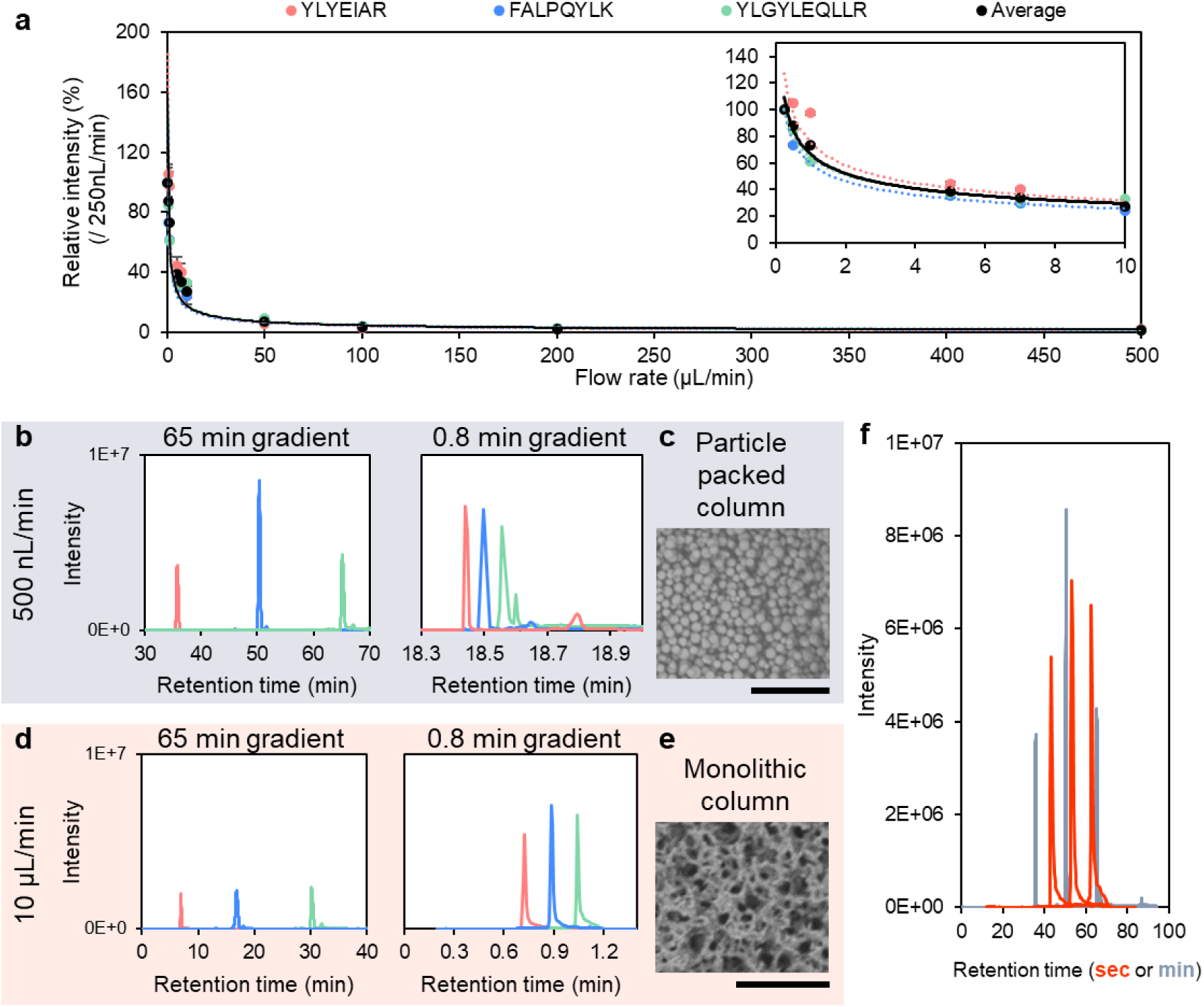
Relationship between electrospray ionization efficiency, flow rate and gradient time. (a) Dependence of flow rate on ESI efficiencies for three synthetic peptides. The x-axis shows flow rate and the y-axis shows intensity relative to intensity at 250 nL/min. The inset shows the expanded range of 250 nL /min ∼ 10 µL/min. Regression curves were drawn by power approximation. (b) Extracted ion chromatograms (XICs) of three synthetic peptides for 65 minute and 0.8 minute gradient elutions at 500 nL/min. (c) SEM image of a particle-packed column (ReproSil-Pur 120 C18-AQ, 1.9 µm particle, 100 µm diameter, 150 mm length) used for (b). Scale bar indicates 20 µm. (d) XICs of three synthetic peptides for 65 minute and 0.8 minute gradient separations at 10 µL/min. (e) SEM image of a monolithic silica column (MonoCap C18 High Resolution 2000, 100 µm diameter, 150 mm length) used for (d). Scale bar indicates 20 µm. (f) Overlaid XICs of three synthetic peptides separated with a 65 min gradient of 500 nL/min (gray) and a 48 sec (0.8 min) gradient of 10 µL/min (red).

### Relationship between flow rate and LC gradient time

Next, the effects of the steep LC gradient elution on speed and sensitivity were examined under nano-flow and capillary-flow conditions. In a typical nano-flow gradient analysis of 65 min, the three synthetic peptides were observed as three individual peaks (Fig. 1b). However, under the 0.8 min steep gradient condition, these peptides no longer eluted (Fig. S1a,b). When the gradient was extended to 99%B at the same gradient slope, these peptides eluted and the peak width became narrower than at the 65-minute gradient analysis (Fig. 1b). Although the narrower peak width was expected to result in a higher peak height, its intensity was not much different from that of the 65-minute gradient. In addition, multiple peaks were observed for each peptide. This could be because the gradient time was too short, resulting in a lower volume of eluent relative to the column volume, and all of the peptides retained on the column could not be eluted. Alternatively, the too fast nano-flow gradient analysis may have resulted in poor mixing of mobile phases A and B. The shorter gradient time brought forward the retention time of each synthetic peptide by about 10 minutes, speeding up the measurement to some extent, but the time required for sample loading, column washing, and equilibration remained the same, and the cycle time ended up taking about 50 minutes. Therefore, in nano-flow LC/MS/MS, it proved difficult to construct a high-speed LC/MS/MS system simply by shortening the gradient time.

Combining a capillary- to micro-flow (1 - 50 µL/min) with a 15-25 cm long LC column (Fig. 1c) with a narrow inner diameter and sub-2 micron size packing material, which is common in shotgun proteomics, was difficult due to the pressure limit of the LC system. The use of a wide-bore column and large particles for low pressure would compromise sensitivity and LC resolution. Therefore, we focused on monolithic silica columns^29, 30^ with wide through-pore (Fig. 1e). When measuring at a flow rate of 500 nL/min using a monolithic silica column with an inner diameter of 100 µm and a length of 2000 mm, the pressure is at most 200 bar. In fact, using a C18 monolithic silica column with an inner diameter of 100 μm and a flow rate of 10 μL/min, we found that the pressure was about 200 bar at room temperature, which is within the pressure tolerance of standard HPLC instruments. Using this monolithic column, synthetic peptides were measured with capillary-flow gradients of 65 min and 0.8 min (Figure 1d); peptide retention time was 30 min earlier in the 65 min gradient analysis than in the 0.5 μL/min system due to the higher flow rate. However, as expected from the ionization efficiency experiments, the peptide peak height was reduced by about 25∼50%. When the gradient time was reduced from 65 min to 0.8 min at the same 10 μL/min flow rate, peak sharpening was observed, with the peak height increased by about 3∼4 times (Figure 1d). Interestingly, comparing the two results under the gradient analysis at 10 μL/min for 0.8 min and 500 nL/min for 65 min, the peak heights were almost equivalent (Fig. 1f). In other words, the combination of capillary-flow and subminute gradient suggested that capillary-flow LC/MS/MS can be significantly accelerated while maintaining sensitivity comparable to that of commonly used nano-flow methods in shotgun proteomics.

### Establishment of machine-gun proteomics

Optimization of flow rate under capillary-flow conditions was performed in ddaPASEF^26^ mode using 500 ng of HeLa digest. As a result, the peak width at half height of the commonly identified 336 peptide peaks became smaller as the flow rate increased (Fig. 2a). The median intensity of the identified peptide peaks and the number of identified peptide-spectrum matches (PSMs) reached a maximum at 7 µL/min, followed by 10 µL/min (Fig. 2b,c).

**Figure 2.**
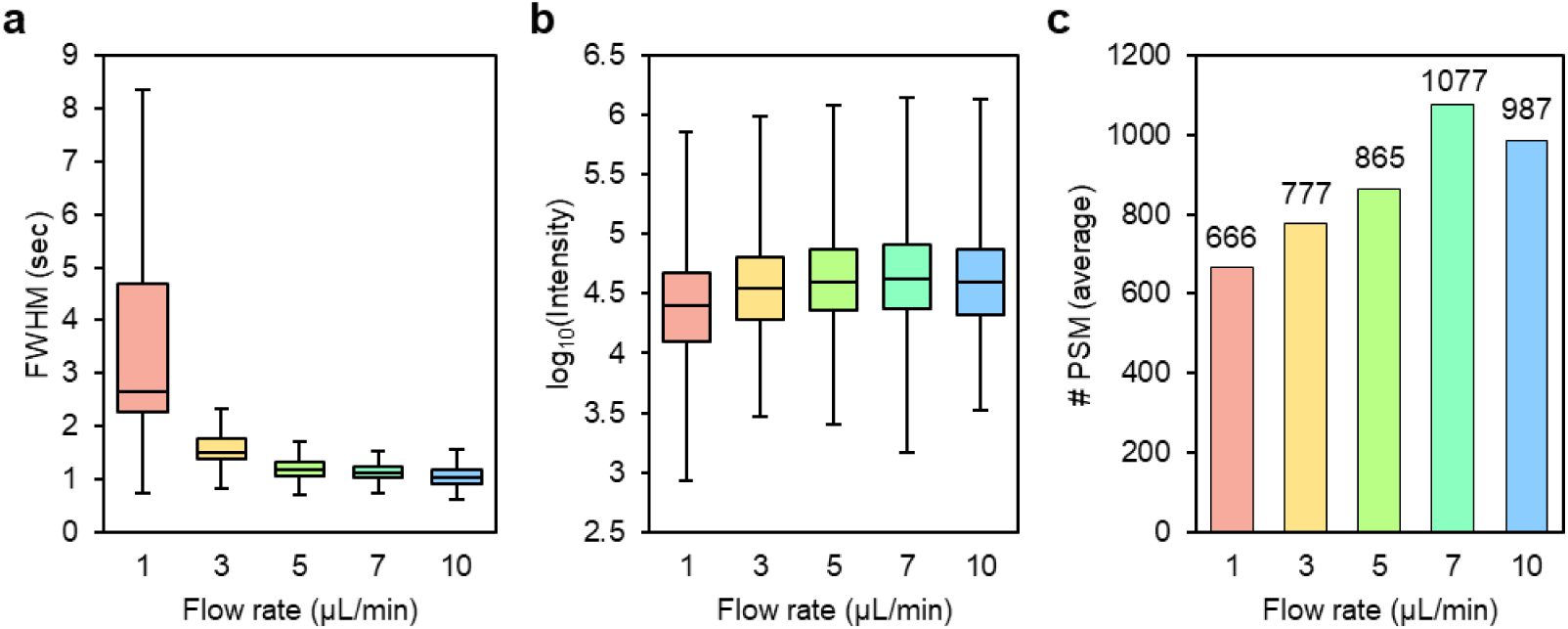
Optimization of LC conditions of machine-gun proteomics. (a) - (c) Box plots of peptide half-widths (a), box plots of logarithm of intensity (b), and number of PSMs (c) when gradient time is set to 1 min and flow rate is varied to 1, 3, 5, 7, and 10 µL/min. The sample used 500 ng of HeLa cell digest.

We then attempted to minimize the dead volume of the LC system. Although it is possible to shorten the LC cycle time by using two trap columns in parallel and injecting the next sample into the trap column during the gradient measurement with another trap column^22^, it is expected to complicate the system and sample loss. In this study, a simple and robust system with as few fittings as possible was selected. Behind the gradient pump and gradient mixer, there is a sample loop connected to a 6-port injection valve, and the sample solution is loaded directly by syringe. After switching the valve, the gradient eluent flows through the sample loop to the column. This system is expected to minimize carryover since the sample loop is flushed throughout the LC measurement time. To reduce dead volume, LC tubing with the smallest diameter must be used, considering that too small-bore tubing increases the system pressure.

In this study, we selected a fused silica capillary with an inner diameter of 50 µm. To keep the length of this capillary tube as short as possible, we selected LC pumps with the external gradient mixer and placed this mixer as close as possible to the injection valve, minimizing the volume from the LC to the injection valve to 0.29 µL. A 1 µL sample loop was also selected to ensure both injection accuracy and minimum volume. In addition, by keeping the distance between the injection valve and the MS instrument as close as possible, the flow path volume from injection to the column head was reduced to 0.29 µL. These considerations reduced the dead volume of the LC system to 1.6 µL in total.

Next, we investigated sample preparation steps to reduce carryover and LC cycle time. In particular, peptide elution conditions during desalting in StageTip were examined to reduce the wash time of the LC analytical column were investigated (Supplementary Fig. 2a, b). The blank and HeLa measurements were conducted alternately, and the analytical column was washed with 99%B (80% acetonitrile in 0.5% acetic acid) in 0 or 2 minutes. Carryover was calculated from the ratio of the blank peak area to the HeLa peak area (Supplementary Fig. 2c). Carryover was 4.2 - 5.2 % when the acetonitrile concentration of the StageTip elution was 80 %, and carryover increased when the washing time was 0 min (Supplementary Fig. 2c). However, by reducing the acetonitrile concentration for the StageTip elution to 32 %, the carryover was reduced to 1.6 - 1.7 %, and under these conditions, no increase in carryover was observed even when the washing time was 0 min (Supplementary Fig. 2c). As mentioned above, the LC system employed in this study has the disadvantage of unavoidable gradient delay due to the volume of the sample loop as the gradient eluent passes through the sample loop, while this system has the advantage of carryover reduction. By optimizing elution conditions in StageTip, washing the sample loop with gradient eluent, and setting a minimum dead volume of 1.6 μL, this system successfully minimized carryover while minimizing the impact on the LC cycle time.

The next step was to examine the column equilibration time. When the column equilibration time was set to 1 min and the LC cycle time to 2 min, peptides eluted up to 1.6 min (Supplementary Fig. 3a). Therefore, we examined whether the reproducibility of retention time could be maintained even if the equilibration time was reduced to 0.6 min in the LC gradient program. First, the column was equilibrated with 5% B for a sufficient time as a blank, and three consecutive measurements were performed with a gradient time of 1 min and an equilibration time of 0.6 min. In the first and second measurements, a slight disturbance in reproducibility was observed for peptides with early retention times, but the retention times in the second and third measurements were unchanged (Supplementary Fig. 3b-e). Therefore, we determined that an equilibration time of 0.6 min was sufficient although not completely back to the initial state. Based on these results, we succeeded in establishing the 720 SPD system with an LC cycle time of 2 min, which we named machine-gun proteomics.

To confirm the reproducibility of repeat injections, 500 ng of HeLa digest was measured 100 consecutive times in ddaPASEF mode with MS duty cycle time set to 400 msec. The average number of identified peptides was 677 ± 41, and the coefficient of variation (CV) values of retention time and peak height were 1.9 - 2.5 % and 8.4 - 10.9 %, respectively, for the three commonly identified peptides over the 100 measurements, which is comparable to that of general nano-flow LC/MS/MS.

### Benchmarks of machine-gun proteomics

We evaluated the quantitative performance of the 720 SPD system using 36 synthetic phosphotyrosine (pY) containing peptides. These 36 synthetic pY peptides derived from trypsin-digested human tyrosine kinases were prepared at various concentrations (10, 25, 50, 75, 100, and 150 fmol/μL for each) as mixtures, and measured in triplicates. In addition, a phosphopeptide-enriched sample from 50 µg of HeLa digest (approximately 250 ng) was spiked to each mixture as the background matrix to determine if quantification was possible under more complex conditions. The synthetic pY peptides were labeled by reductive dimethylation with light isotopes (dimethyl light label) while the matrix phosphopeptides were labeled with dimethyl heavy isotopes (Fig. 3a). Single-stage MS without MS/MS measurements were performed using the 720 SPD method with the MS duty cycle time of 100 msec. The heat map shows the results of the linearity experiments between sample concentration and peak area for the 36 pY peptides (Fig. 3b). For samples without matrix, high linearity was observed for most of the pY peptides (median R^2^ = 0.990). In addition, the addition of matrices did not significantly impair linearity (median R^2^ = 0.988). This suggests that our machine-gun proteomics method can accurately quantify peptides even in complex samples. Regarding the repeatability, no concentration dependence was observed, with a median CV of 5 - 15% for each concentration (Supplementary Fig. S5). The TICs and XICs of five representative synthetic pY peptides were then examined in the presence of the matrix. Three of them were completely separated without interference, while the remaining two overlapped with each other (Fig. 3c). These two peptides have the same sequence but different phosphorylation localization (Ephrin type-A receptor 2 (EphA2) 918-935, 921Y or 930Y). Eph receptor-mediated signaling is known to regulate key cellular functions such as cell migration, axon guidance, and angiogenesis under physiological and pathological conditions, including cancer^31^. It was reported that phosphorylation at 921Y and 930Y enables differential binding to the Src homology 2 domain of the adaptor protein Grb7, leading to distinct functional outcomes^32^. Also, receptor-type tyrosine-protein phosphatase F (PTPRF) specifically dephosphorylates 930pY on EphA2 to control its association with Nck1 and subsequently, cell migration^33^. Thus, these two phosphorylation sites must be quantified separately. Since only the phosphorylation localization differed, the *m/z* values were identical, and without ion mobility filtering, the separation based on retention time alone was insufficient and the peak boundaries were ambiguous (Fig. 3c). However, with the addition of ion mobility separation, the two peptides could be extracted as distinct peaks (Fig. 3c,d). The linearity was also improved by adding the ion mobility information (Fig. 3e,f). In other words, it was found that the separation by ion mobility compensated for the deterioration of LC separation due to short gradient time in this method. Note that endogenous 921Y and 930Y phosphopeptides were detected with heavy labels in this assay, but their content was less than 10 fmol/μL, which is outside the range of the calibration curve.

**Figure 3.**
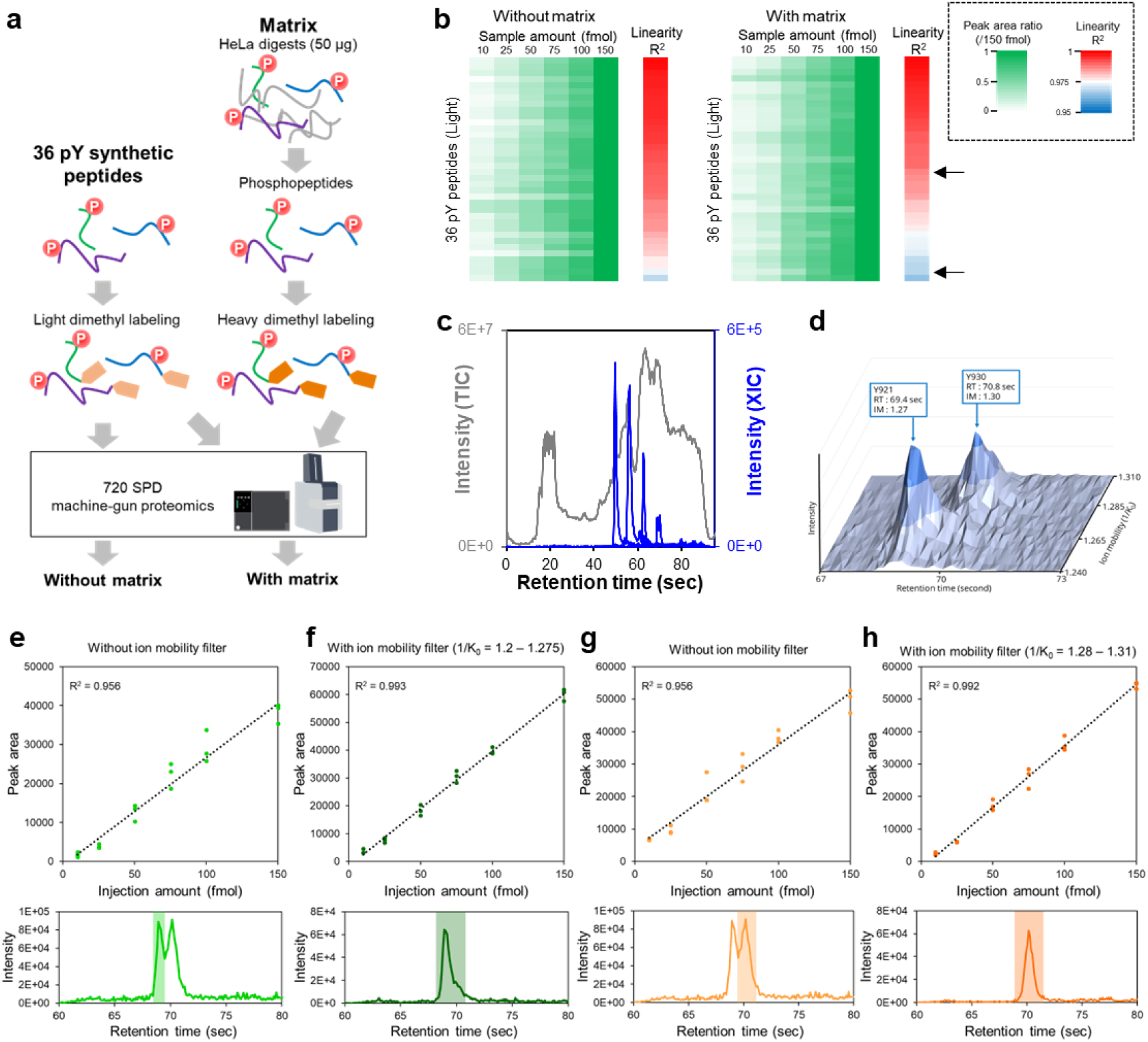
Linearity assessment in quantitative machine-gun proteomics using 36 phosphotyrosine synthetic peptides. (a) Experimental design for quantitative analysis of 36 pY peptides by machine-gun proteomics. (b) Linearity evaluation overview. The green to white heatmap shows the peak area ratio relative to the result of spiking 150 fmol of pY synthetic peptide. Red, white, and blue bars indicate R-squared values of linearity. The two arrows point to EphA2 pY921 (bottom) and pY930 (top), respectively. (c) TIC in the “with matrix” experiment (gray) and the overlaid XICs of the five pY peptides (blue). (d) Two-dimensional separation of EphA2 921pY and 930pY peptides by retention time and ion mobility. (e-h) Top: scatter plots showing the linearity between spike amount and peak area for pY synthetic peptides. Bottom: XICs of synthetic peptides. (e) and (g) show the results for EphA2 pY921 and pY930, respectively, without ion mobility filter. (f) and (h) show the results for EphA2 pY921 and pY930, respectively, with ion mobility filter.

We then evaluated the performance of machine-gun proteomics using diaPASEF^34^ with injection amounts varying from 10 ng to 1000 ng of HeLa digests (triplicate injections). The MS acquisition was performed with one MS1 scan followed by four PASEF scans. The TIMS ramp time was set to 100 msec and the MS duty cycle time was 500 msec (See details in Materials and Methods). To construct the spectral library, the 48 fractions were pre-fractionated by reversed-phase HPLC, concatenated into 12 fractions, and each was subjected to LC/MS/MS measurements in DDA mode under the same conditions as for machine-gun proteomics. As a result, we were able to identify up to 20,000+ precursor ions and 3,500+ protein groups. (Fig. 4a). Furthermore, even when using samples as small as 10 ng, about 2,000 protein groups were identified (Fig. 4a). This confirmed that our method can simultaneously achieve high throughput, high sensitivity, and depth of analysis. However, the 720 SPD showed a decrease in the identification number at >1000 ng. This may be due to the shortening of the gradient, which resulted in too many peptides eluting at one time, leading to saturation. In fact, the peak width tended to become wider as the sample amount was increased, suggesting that this may have been caused by a deterioration in peak capacity (Fig. 4b). The reproducibility in quantitation was also examined and found to be 11-18%, which is comparable to DDA quantitation (Fig. 4c). To further increase throughput, a 1000 SPD method with a steeper gradient of 0.8 min was also launched and its performance was evaluated. As a result, higher identification values than the 720 SPD method were obtained for samples from 10 ng to 100 ng, and the 2145 and 3139 protein groups could be identified from 10 ng and 100 ng, respectively. As for quantitative reproducibility, the median CV was less than 20%, and as with the 720 SPD method, performance tended to deteriorate gradually as the sample a ount exceeded 250 ng.

**Figure 4.**
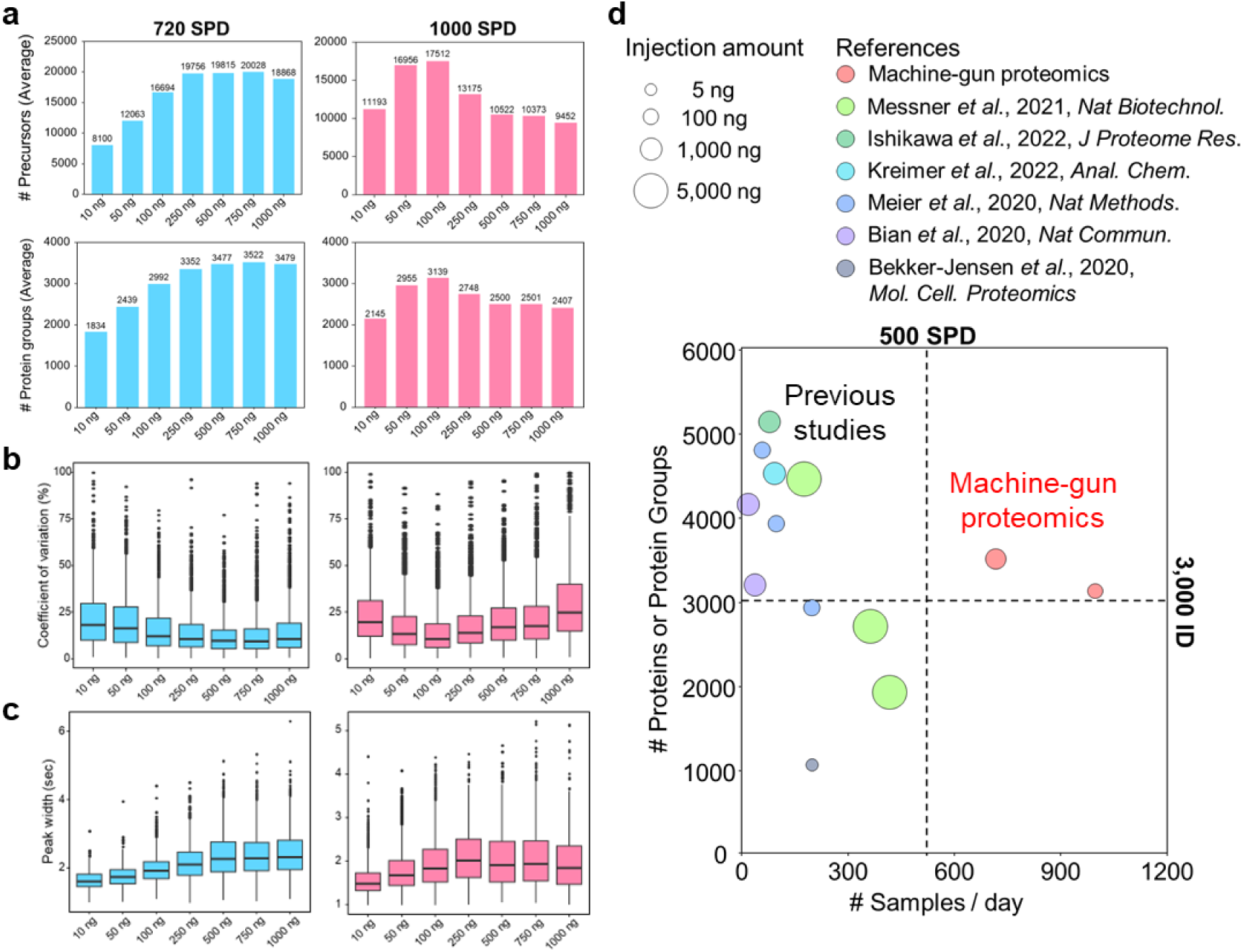
Benchmarks of machine-gun proteomics with HeLa dilution series. (a) Bar graphs of average of the identification number of the precursor (top) and protein groups (bottom). (b) Boxplot of repeatability in protein quantification by DIA analysis (c) Boxplot of peak width of commonly identified precursor ions. (d) Bubble chart comparing previous studies with this study. X-axis indicates the number of samples that can be measured per day, y-axis indicates the number of identified protein or protein group, and bubble size indicates the sample injection amount of human cell line digests.

Next, we compared our methods with previously reported literatures related to high throughput proteomics (Fig. 4d). Our methods were able to measure more than double the number of SPD compared to the fastest method reported so far, with the same protein identification number. The sample amount was also smaller (Messner et al., 5,000 ng, our study, 100 or 750 ng), indicating that high sensitivity was achieved. In summary, our method did not sacrifice any of the three aspects of LC/MS/MS throughput, sensitivity, or depth of analysis.

#### Spatially resolved proteome analysis of mouse brain

We applied our machine-gun proteomics platform to spatial proteomics, which requires high sensitivity and high throughput measurement. A total of 96 sections of ∼440 μm square were cut from one mouse brain slice using LMD, and each section was collected into each well on a 96-well plate. For a total of 96 samples, proteins were extracted, digested, and measured by the 1000 SPD machine-gun proteomics method. An average of more than 500 protein groups was identified in a total of 96 sections cut from each LMD (Fig. 5b). The number of identified proteins was lower than in the benchmark results (Fig. 4a), but this can be due to losses in sample preparation, in addition to the limited number of expressed proteins in the mouse brain. GO enrichment analysis of the identified proteins using DAVID2021 revealed significant enrichment of GO terms for brain-related cellular components such as myelin sheaths and synapses (Fig. 5c). Next, we examined the spatial expression information of the identified proteins. No significant differences by spatial position were observed for precursor ions, peptides, and number of protein groups (Fig. 5b). This was also true for the spatial distribution of trypsin-derived peptides added during sample preparation (Fig. 5d). On the other hand, myelin basic protein (Mbp), myelin proteolipid protein (Plp1), and calcium/calmodulin-dependent protein kinase type II subunit alpha (Camk2a), unlike trypsin, showed a certain distribution along the structure of mouse brain sections (Fig. 5e-h). The pattern of spatial distribution of these quantitative values was confirmed repeatedly for different peptides derived from the same protein. Mbp and Plp1, which are known to be well expressed in myelin, were confirmed to have almost the same spatial distribution, although they are different proteins (Fig. 5g, h). These results suggest that our machine-gun proteomics approach can accurately quantify spatial protein expression patterns.

**Figure 5.**
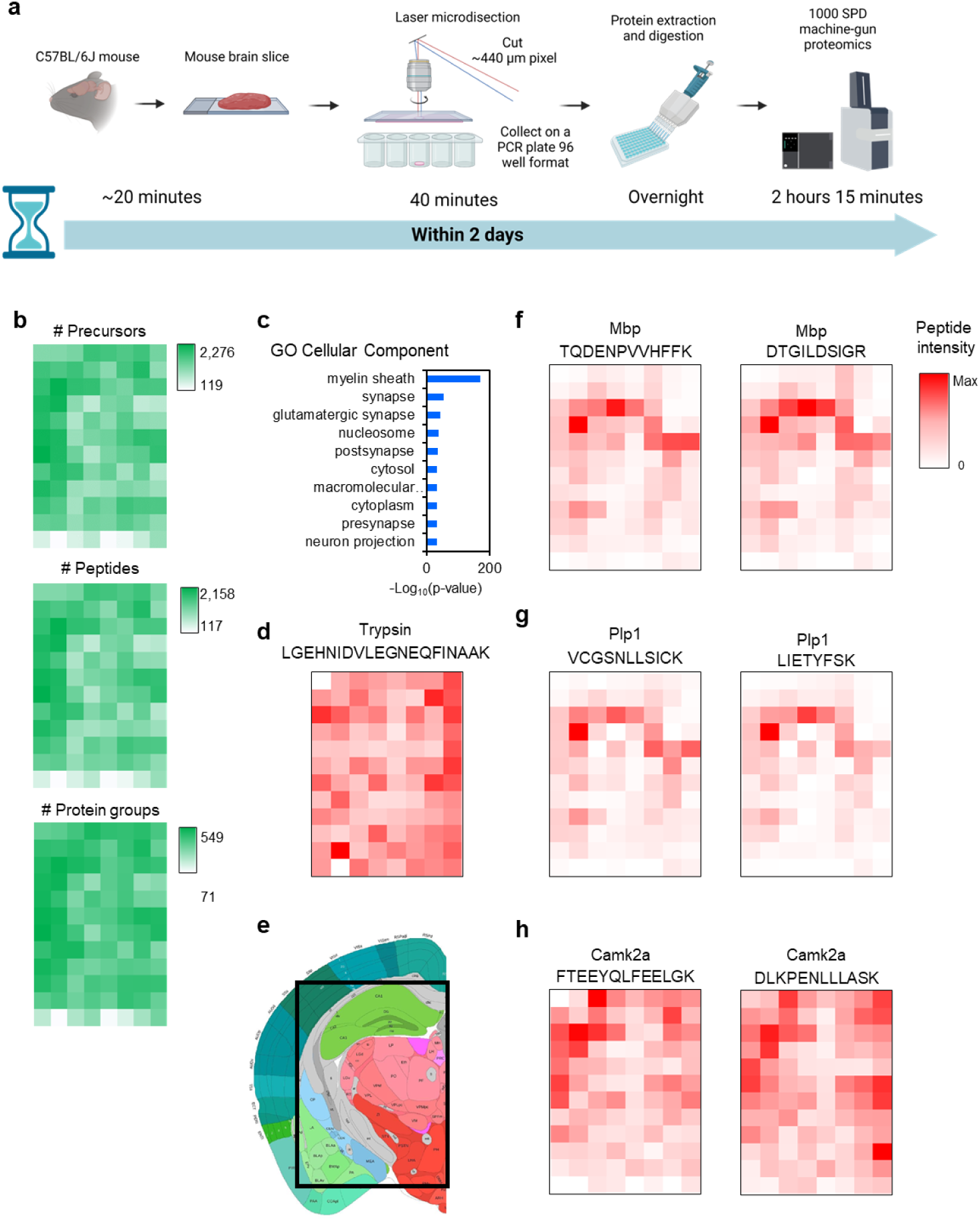
Spatial proteomics of mouse brain by machine-gun proteomics. (a) Experimental workflow of spatial proteomics and its required time. This figure was created with BioRender.com (b) Bar plot of the number of identified precursors (top), peptides (middle), and protein groups (bottom). (c) Bar plot of GO enrichment analysis. (d) Heat maps of the spatial distribution of signal intensities of trypsin. (e) Mouse brain section site used in this experiment. Data are from the Allen Mouse Brain Atlas^35–38^ (mouse.brain-map.org). (f) - (h) Heat maps of the spatial distribution of signal intensities of myelin basic protein (Mbp), myelin proteolipid protein (Plp1), and calcium/calmodulin-dependent protein kinase type II subunit alpha (Camk2a) derived peptides.

## Discussion

We have developed machine-gun proteomics, the highest-speed proteome measurement system without sacrificing sensitivity and the depth of analysis. While previous high-speed methods have achieved high robustness with high LC flow rates of several hundred µL/min or so, their sensitivity has been reduced. Direct infusion MS without LC was also reported to achieve high speed, but it was difficult to overcome the high complexity of proteome samples, and the protein identification number was limited to about 500^25^. So far, attempts have been made to achieve both throughput and sensitivity at high flow rates, such as capillary-flow, but the throughput has been inferior to that of high flow rate systems^21–24^.

In this study, we investigated ESI efficiency in the flow rate range and quantitatively evaluated the relationship between flow rate and ESI efficiency. We also showed that the decrease in sensitivity with increasing flow rate can be compensated for by shortening the gradient time. Based on these basic investigations, we succeeded in achieving both throughput exceeding that of micro-flow LC/MS/MS and sensitivity as high as that of nano-flow. Note that this method is basically a customized version based on a commercial device and can be reproduced in other laboratories. The key point is that the gradient mixer was placed outside the LC system, and the MS ion source with an ESI emitter-integrated LC analytical column, injection valve, and gradient mixer were placed as close as possible to minimize gradient delay time. In addition, the monolithic column selected for this study is expected to be highly robust, as it does not clog even when measuring large volumes of coarse clinical samples^39^. In fact, we routinely perform several hundred consecutive measurements, and the frequency of column replacement is extremely low.

The duty cycle time for the three MS data acquisition modes used in this study was 100 msec for the pY peptide analysis, 400 msec for 100 replicate measurements by DDA, and 500 msec for the HeLa dilution series experiment by DIA. Since the average peak width at half height of the LC is about 1 sec, there was concern that the number of data points per peak in DDA and DIA modes, in particular, would worsen the reproducibility of retention times and quantitative values. However, the CV values of retention time for commonly identified peptides was comparable to that of typical nano-flow LC/MS/MS, averaging about 3.4% over 100 repeat measurements for DDA. The similar performance was observed for HeLa dilution series experiments using the DIA mode, i.e., the quantification precision of the identified proteins was approximately 11 - 18% for 720 SPD and 12 - 24% for 1000 SPD, which is sufficient for quantitative proteomics experiments.

For spatial proteome analysis of mouse brain, Piehowski et al. performed LC/MS/MS analysis of 100 µm square sections and MS imaging of 2,000 proteins over 3 hours per section^40^. In contrast, using our method, the LC/MS/MS analysis of one section took only 1.4 min and the total LC/MS/MS analysis for 96 sections was 2 hour 15 min, which is less than one section analysis by Piehowski et al.^40^. In other words, we successfully accelerated the measurement more than 100-times, although the protein digestion step took overnight.

The spatial resolution of the proteomics in this study is low at 400 μm square. However, the sample preparation method used in this study is the same as that used for general large-amount sample preparation, so further improvement can easily be expected by using the high-sensitivity sample preparation methods that have been reported in various fields, especially single cell analysis, in recent years^9, 41^.

Although spatial proteomics is shown as an application example, our machine-gun method can be applied to a wide range of fields. For example, applications to biomarker discovery at the proteome level and personalized medicine are expected in the future. Another advantage is that the method is not only fast but also highly sensitive, so it can be challenged regardless of sample amounts.

In conclusion, our “machine gun proteomics” system is a major step forward in fast and sensitive LC/MS/MS for comprehensive proteome analysis. By optimizing flow rates and incorporating ion mobility-based separations, we have overcome previous limitations and achieved high throughput, high sensitivity, and high depth analysis. This breakthrough has important implications for biomarker discovery, spatial proteomics, and other areas of proteomics research, facilitating a deeper understanding of complex biological processes.

## Materials and Methods

### Materials

Sodium deoxycholate (SDC), sodium *N*-lauroylsarcosinate (SLS), dithiothreitol (DTT), iodoacetamide (IAA), ammonium bicarbonate, MS-grade Lysyl endopeptidase® (Lys-C), ethyl acetate, acetonitrile (ACN), acetic acid (AA) and trifluoroacetic acid (TFA) were obtained from FUJIFILM Wako (Osaka, Japan). Modified trypsin was obtained from Promega (Madison, WI, USA). Empore disks were from GL Sciences (Tokyo, Japan). Water was purified with a Milli-Q system Merck (Darmstadt, Germany).

### Preparation of mouse brain tissue

The brain was freshly isolated from C57BL/6J male mice at 8 weeks of age and snapped frozen with liquid nitrogen. Tissue was then coronally sectioned into 10 μm thickness with Leica CM3050 S cryostat and placed on the DIRECTOR™ slides (AMR, Tokyo, Japan).

### Laser microdissection

Laser microdissection and sample collection were performed using a Leica LMD7 system (Leica Microsystems GmbH). A frozen section of mouse brain (coronal) was divided into 96 pixels (an area of 191,014 µm^2^ per pixel) at 2.5x objective by auto detection mode software module. The tissue section was dissected with UV laser scanning and samples were collected by gravity drop into a 96-well plate that contained the PTS buffer with 10 mM tris(2- carboxyethyl)phosphine (TCEP) and 40 mM chloroacetamide (CAA).

### Sample preparation of HeLa dilution series for benchmarking machine-gun proteomics

HeLa cells were grown in DMEM until 80% confluence in 10 cm diameter dishes. Cells were washed twice with cold phosphate-buffered saline (PBS) and collected by spinning at 500 g for five min at 4 °C and the supernatant was discarded. Collected cells were lysed and digested according to the phase transfer surfactant (PTS) protocol^42^ with some modifications as follows. After cell lysis with PTS buffer (10 mM SDC, 10 mM SLS, 100 mM Tris-HCl (pH 9) and protease and phosphatase inhibitors), the tubes were heated at 95°C in a thermal cycler, and ultrasonicated for 20 minutes at ice-cold temperature. The amount of protein was quantified with a BCA protein assay kit. The denatured proteins were reduced by 10 mM DTT at room temperature for 30 minutes and alkylated with 50 mM IAA in the dark at room temperature for 30 minutes. The proteins were 5-fold diluted with 50 mM ammonium bicarbonate. LysC were added at the ratio enzymes: proteins = 1:100 and incubated several hours at room temperature, after that trypsin were added at the ratio enzymes: proteins = 1:100 and incubated overnight at 37°C. The next day, an equal volume of ethyl acetate was added to remove surfactant, and digestion was stopped by adding 5 µL of 20 % TFA. The mixture was shaken for 2 minutes and centrifuged at 15,800 g for 2 minutes. Then, the aqueous phase was collected. Digested samples were desalted using StageTips^43, 44^ with SDB-XC Empore disk membrane.

### Phosphopeptides enrichment and dimethyl labeling for evaluation of quantitative linearity using pY peptides

The HeLa-derived phosphopeptides were enriched by hydroxy acid-modified metal oxide chromatography (HAMMOC) with elution by 0.5% piperidine^45, 46^. After enrichment of phosphopeptides, samples were desalted by StageTips and resuspended for capillary-flow LC/MS/MS analyses.

Stable isotope dimethyl labeling was carried out as previously described^47^ with some modification as follows. Briefly, the HeLa-derived tryptic phosphopeptides or synthetic phosphopeptides were dissolved in 1000 µL of 100 mM TEAB. Then, the peptides were mixed with 40 µL of 4% ^13^CD_2_O (HeLa-derived tryptic peptides) or ^12^CH_2_O (synthetic peptides), and 40 µL of 0.6 M sodium cyanoborohydride was added. The mixture was stirred for 60 min at room temperature. The reaction was stopped by adding 160 µL of 1% ammonium hydroxide and then samples were mixed with 620 µL of 10% TFA. The labeled peptides were desalted by using SDB-XC StageTips^43, 44^. After labeling, synthetic peptides were spiked with phosphorylated peptides from HeLa cells. When spiking, the concentration was 10, 25, 50, 75, 100, and 150 fmol for phosphopeptides from 50 μg of HeLa cell digests.

### Sample preparation for spatially resolved proteomics on mouse brain

Tissue samples were collected by automated LMD into 96-well plates with 20 µL of 12 mM SDC, 12 mM SLS, 100 mM Tris-HCl (pH 9). Dissected mouse brain tissues were digested according to the phase transfer surfactant (PTS) protocol^42^ with some modifications as follows. After tissue collection, plates were heated at 95 °C in a thermal cycler, and ultrasonicated for 20 minutes at ice-cold temperature. The denatured proteins were reduced by 10 mM DTT at room temperature for 30 minutes and alkylated with 50 mM IAA in the dark at room temperature for 30 minutes. The proteins were 5-fold diluted with 50 mM ammonium bicarbonate. 1 μL of trypsin was added and incubated overnight at 37 °C. The next day, an equal volume of ethyl acetate (100 µL) was added to remove surfactant, and digestion was stopped by adding 5 µL of 20 % TFA. The mixture was shaken for 2 minutes and centrifuged at 15,800 g for 2 minutes. Then, the aqueous phase was collected. For all samples, digests were desalted with StageTips^43, 44^ with SDB-XC Empore disk membrane.

### Off-line high pH reversed phase peptide fractionation for spectral library generation

For 720 SPD HeLa benchmarks, the Shimadzu LC-30AD system operating a L-column3 C18 150 mm × 2.1 mm, 3 µm column was used to fractionate peptides at a flow rate of 200 μL/min. Mobile phase A was 0.1% triethylamine (TEA), mobile phase B was 0.1 %TEA, 80 %ACN. The 200 μg HeLa digests were separated by a linear gradient of 2 -35%B in 65 minutes. Fractions were collected every 1.33 minutes and fractions were collected into a 96 deep well plate. For 1000 SPD HeLa benchmarks, Shimadzu Nexera Mikros LC pump operating a Waters nanoEase™ M/Z Peptide 300 μm × 100 mm, BEH C18, 130 Å 1.7 μm column was used to fractionate peptides at a flow rate of 3 μL/min. Mobile phase A was 0.1% triethylamine (TEA), mobile phase B was 0.1 %TEA, 80 %ACN. The 50 μg HeLa digests were separated by a linear gradient of 2 - 35%B in 65 minutes. Fractions were collected every 1.33 minutes and fractions were collected into a 96 well plate.

The collected 48 fractions were concatenated into 12 fractions by adding fraction 13 to fraction 1, fraction 14 to fraction 2 and so forth. Peptide fractions were dried in a SpeedVac then redissolved in 4% ACN, 0.5% TFA and measured by LC/MS/MS.

### Direct infusion mass spectrometry

Flow injection analysis was performed with an syringe pump 11 Elite (HARVARD Apparatus) connected to a timsTOF Pro 2 mass spectrometer (Bruker) with an electrospray ion source (Bruker). 3 synthetic peptides were dissolved in 10% ACN, 0.5% AA in each concentration (Supplementary Table 1).

### Nano-flow LC/MS/MS

NanoLC/MS/MS analysis was performed with an Ultimate 3000 RSLCnano pump (Thermo Fisher Scientific) connected to a timsTOF Pro 2 mass spectrometer (Bruker) with a nano-electrospray ion source (CaptiveSpray, Bruker). Mobile phase A: 0.5% AA in water; mobile phase B: 0.5% AA, 80% ACN. 5 µL of samples were injected by an HTC-PAL autosampler (CTC Analytics). Peptides were loaded on a in-house pulled capillary column (1.9 µm ReproSil-Pur 120 C18-AQ (Dr. Mainsch) packed in 150 mm length, 100 µm inner diameter fused silica capillary). Then peptides were separated at 0.5 µL/min using a linear gradient from 5–10%B in 5 minutes, 10-40% B in 60 minutes (0.5% AA and 80% ACN), washed in 99% B for 10 minutes, then equilibrated for 30 minutes in 5% B.

The mass spectra were acquired by data-dependent acquisition mode (ddaPASEF)^26^. TIMS accumulation time and ramp time was set to 100 msec. TIMS scan range was 0 - 1.4 V·s/cm^2^. One cycle was composed of 1 MS scan followed by 10 PASEF MS/MS scans. MS and MS/MS spectra were recorded from *m/z* 100 to 1,700. A polygon filter was applied to not select singly charged ions. The isolation width was set to 2 *m/z* for precursor *m/z* < 700 and 3 *m/z* for precursor *m/z* > 800.

### Setup of capillary-flow LC/MS/MS

A Nexera Mikros (Shimadzu) pump was coupled online to a timsTOF Pro or timsTOF Pro 2 mass spectrometer (Bruker) in this study. We used fused silica capillaries (Polymicro Technologies) for all the connections. The pump outlet was directly connected to the P-875 tee connector or M-540 static mixer (IDEX Health & Science) by a 50 μm ID capillary. A 50 μm ID × 150 mm capillary was used to connect the mixer to the sample injection valve. A home-made 1 μL sample loop (100 μm ID × 127 mm capillary) was used in direct injection mode. The sample loop was kept in line during gradient elution. A 50 μm ID × 150 mm capillary was used to connect the sample injection valve to column inlet. Peptides were loaded onto a 15 cm monolithic silica capillary column (100 μm ID, self-pulled needle using home-made C18 monolithic silica column or MonoCap C18 High Resolution 2000, GL Sciences). Mobile phase A: 0.5% AA in water; mobile phase B: 0.5% AA, 80% ACN. A linear gradient of 5-40% B for 0.8 minute or 1 minute, 5% B in 0.6 minute was used for HeLa (phospho)peptides and 3 synthetic peptides derived from BSA digests. For 65 minutes gradient and 10 µL/min measurements of 3 synthetic peptides derived from BSA digests, the linear gradient of 5-10% B in 5 minutes, 10-40% B in 60 minutes, 99% B for 10 minutes, then 30 minutes in 5% B was used.

The mass spectra were acquired by MS1 mode for phosphopeptides linearity experiments, ddaPASEF^26^ for the relationship between flow rate and gradient length experiments and the flow rate optimization experiments, or diaPASEF^34^ for HeLa dilution series or mouse brain experiments.

For ddaPASEF mode, TIMS accumulation time and ramp time was set to 100 msec. TIMS scan range was 0.6-1.5 V·s/cm^2^ for 65 minutes gradient experiments and 0.7-1.4 V·s/cm^2^ for other 0.8 or 1 minute gradient experiments. One cycle was composed of 1 MS scan followed by 10 PASEF MS/MS scans for 65 minutes gradient experiments and 3PASEF MS/MS scans for other 0.8 or 1 minute gradient experiments. MS and MS/MS spectra were recorded from *m/z* 100 to 1,700. A polygon filter was applied to not select singly charged ions. The isolation width was set to 2 *m/z* for precursor m/z < 700 and 3 *m/z* for precursor m/z > 800.

For diaPASEF mode, TIMS accumulation time and ramp time was set to 100 msec. TIMS scan range was 0.7-1.4 V·s/cm^2^. One cycle was composed of 1 MS scan followed by 4 PASEF MS/MS scans. MS and MS/MS spectra were recorded from *m/z* 100 to 1,700. A polygon filter was applied to not select singly charged ions. The isolation width was set to 36 m/z and 0.11 V·s/cm^2^ ranged from *m/z* 400-1120, 1/K_0_ 0.7-1.4, the total number of isolation window was 20 (Supplementary Table 3). The detailed MS/MS acquisition scheme was shown in Supplementally table.

### Data Analysis and Bioinformatics

For the infusion MS analysis, the MS intensity of synthetic peptides were quantified by Bruker Compass DataAnalysis (version 6.0). For the evaluation of relationship between flow rate and gradient time and the benchmark linearity of phosphopeptides quantification analysis, the peptides were quantified by Skyline (version 22.2.0.351) at MS1 level.

For ddaPASEF measurements, peptides were searched by MaxQuant (version 1.6.10.43) against UniprotKB/Swiss-Prot Human database (2017/10 download) with a precursor mass tolerance of 70 ppm, a fragment ion mass tolerance of 40 ppm, and strict trypsin/P specificity allowing for up to 2 missed cleavages. Carbamidomethyl (C) were set as fixed modifications. Methionine oxidation and N-term acetylation was allowed as a variable modification. False discovery rates at a PSM and protein group level of less than 1% were estimated by searching against a reversed decoy database.

For diaPASEF measurements, peptides and proteins were identified and quantified using Spectronaut version 17 in directDIA mode against the human database (download in 04/2017) or mouse database (download in 12/2022) with pig trypsin sequence (download in 03/2023) downloaded from UniprotKB/Swiss-Prot. The search parameters were used as default settings if there are no mentions.

Digestion mode was set Trypsin/P allowing for up to two missed cleavages. Oxidation (M) and acetylation (protein N-term) were allowed as variable modification, and carbamidomethylation (C) was allowed as fixed modification. False discovery rates at precursor, peptide, and protein level of less than 1 % were estimated by searching against a reversed decoy database.

## Data availability

All MS raw files as well as the peak picking and the result files were deposited at a public repository of ProteomeXchange Consortium (http://proteomecentral.proteomexchange.org) via the jPOST partner repository (https://jpostdb.org) with the data set identifier PXD0XXXXX.

All other data are available from the corresponding author on request.

## Acknowledgements

We wish to thank all members of the Ishihama group for technical assistance and fruitful discussions. Parts of this work were funded by grants from the JST Strategic Basic Research Program CREST (No. 18070870), AMED-CREST program (No. JP18gm1010010), JSPS Grants-in-Aid for Scientific Research No. 17H03605 and 21H02459 and JSPS Grant-in-Aid for Challenging Research (Exploratory) No. 20K21478.

## Author contributions

Y. I. Conceptualization. A. T., R. T. and K. O. set up and optimized the capillary-flow LC system. Yo. I., M. A., K. I., E. K., K. O., and Y. I. designed experiments. A. T., R. T., and I. M. performed experiments. A. T., R. T. analyzed data. A. T. and Y. I. writing - original draft. M. A., K. I., E. K., K. O. writing - review & editing.

## Additional information

Supplementary information is available for this paper at …..

## Supplementary information

**Supplementary Figure S1.**
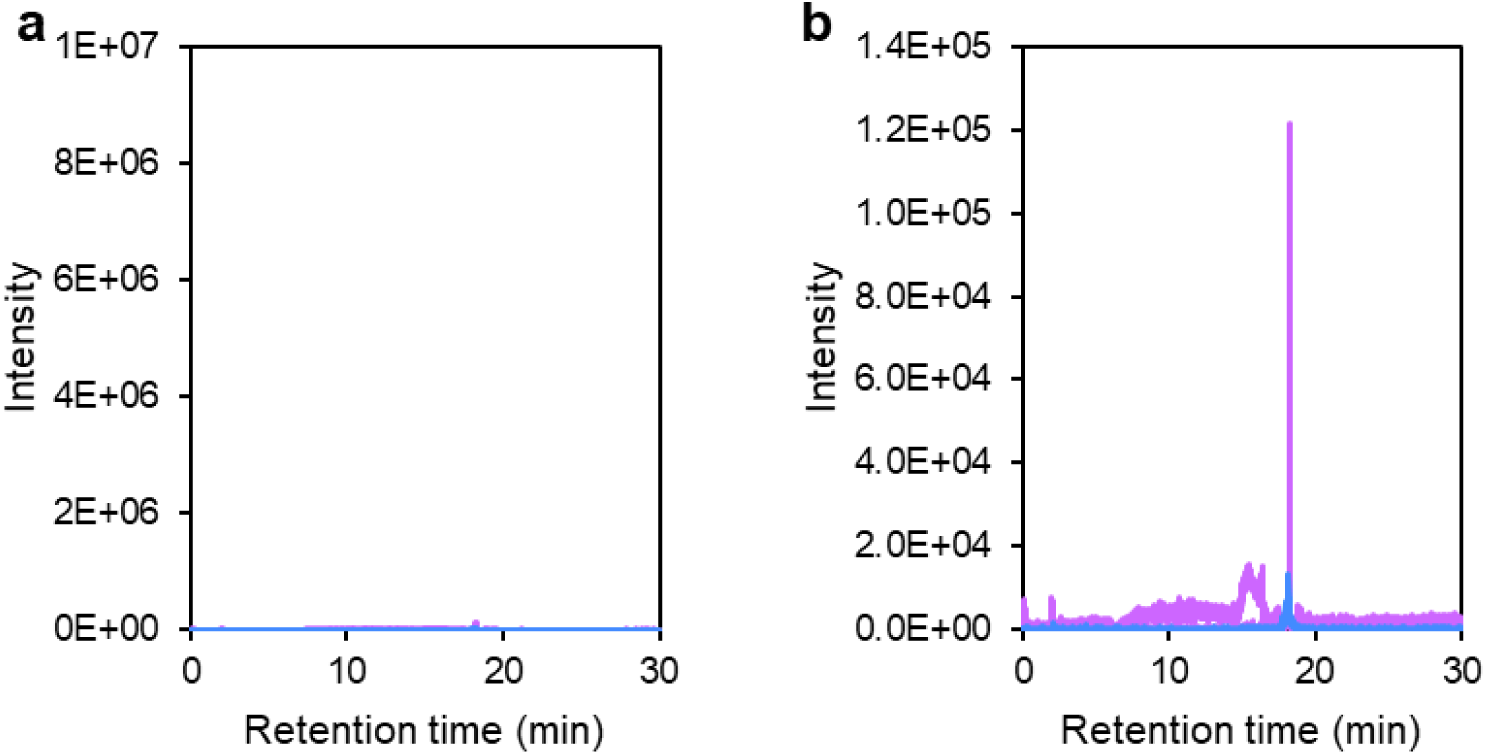
Relationship between electrospray ionization efficiency, flow rate and gradient time. (a), (b) Extracted ion chromatograms (XICs) for 0.8 minute gradient elution at 500 nL/min. The range of the y-axis in (a) is the same as in Fig. 1b and d, while the y-axis in (b) shows the enlarged version.

**Supplementary Figure S2.**
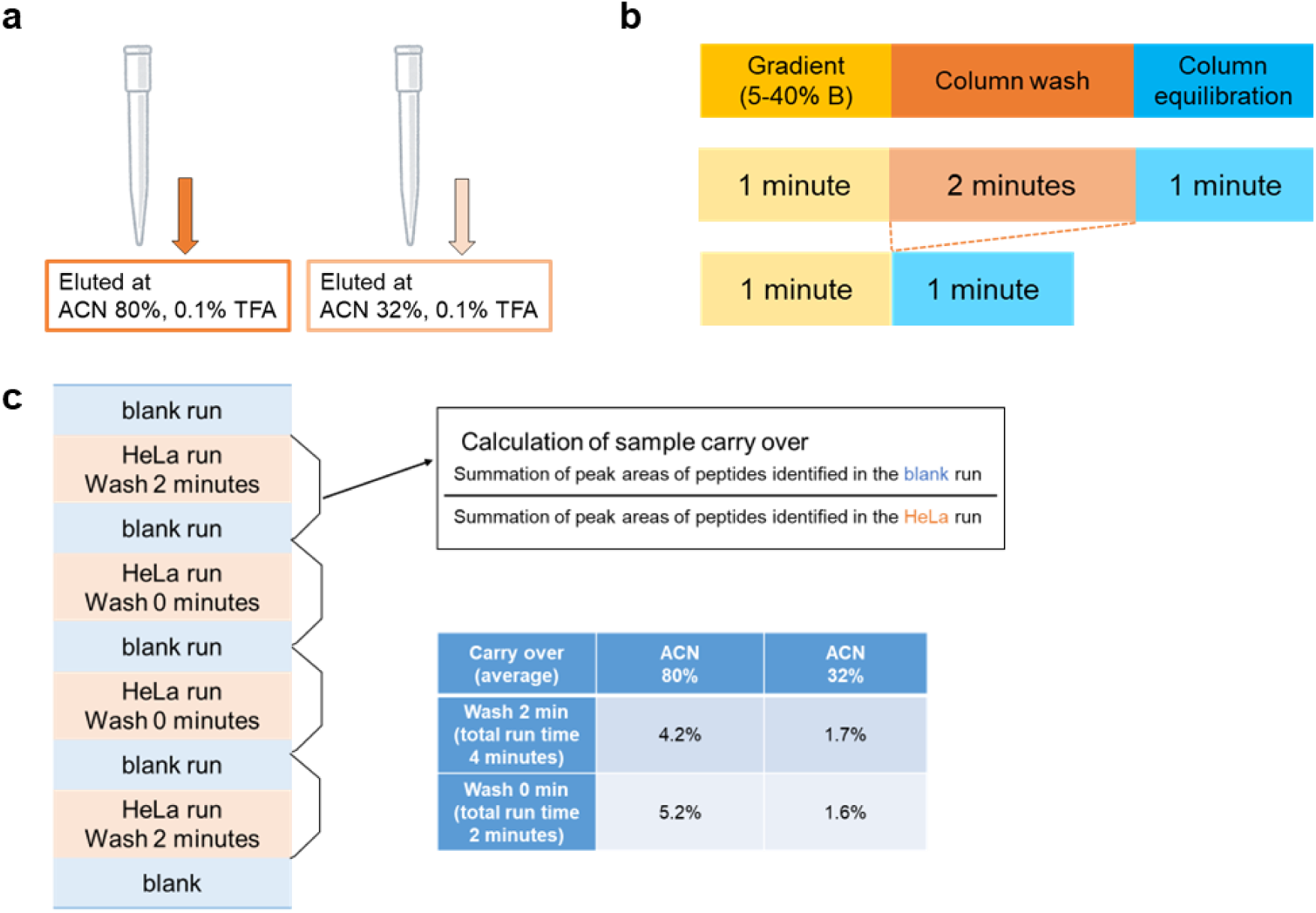
Investigation of LC column wash time. (a) Sample elution conditions during desalting in sample preparation. (b) LC gradient program. (c) Order of LC/MS/MS measurements, method of calculating carryover, and the ratio of sample carry over.

**Supplementary Figure S3.**
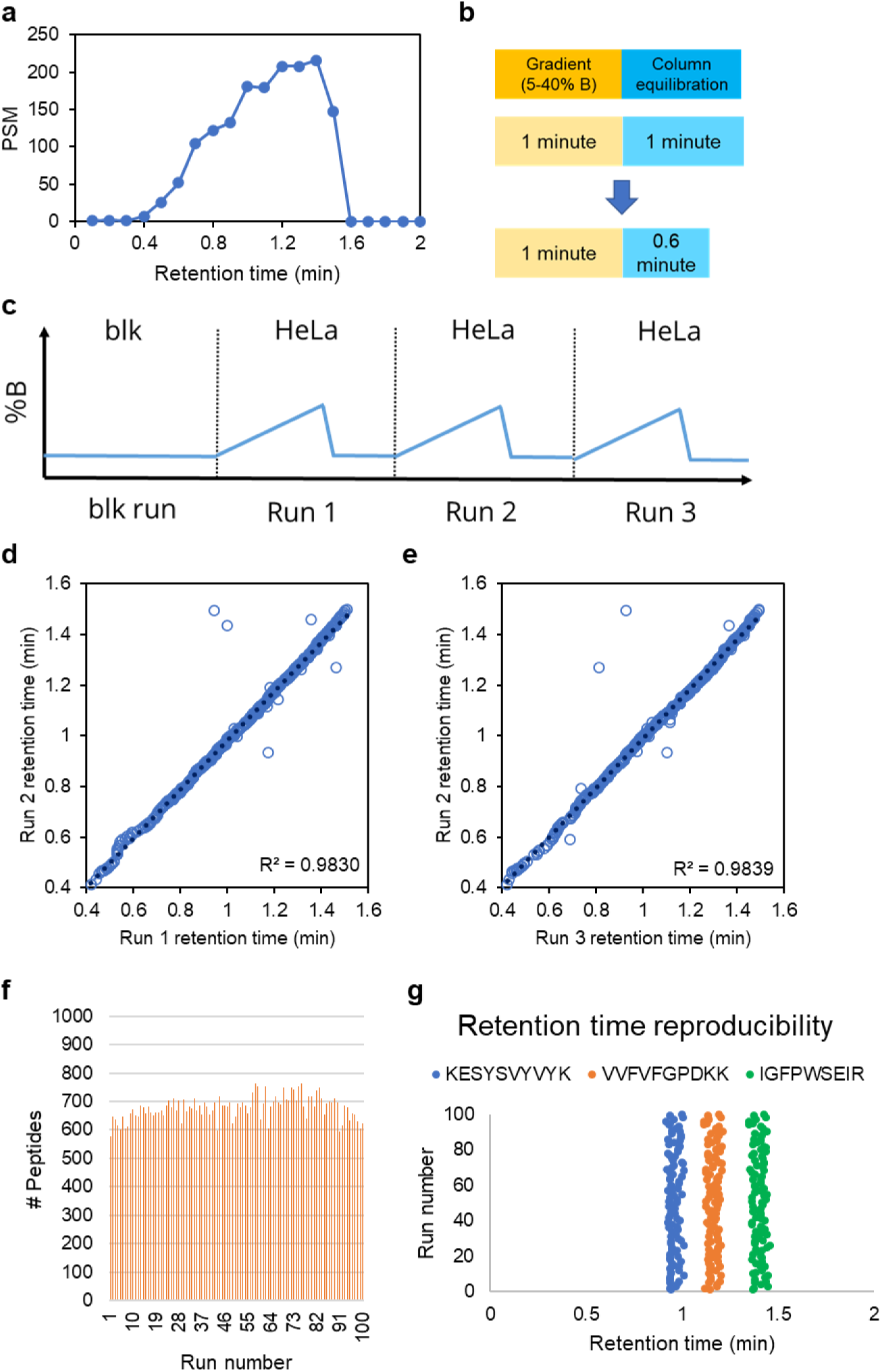
Investigation of LC column equilibration time and robustness of machine-gun proteomics. (a) Retention time distribution of PSM at 7 µL/min, 1 min gradient LC/MS/MS. (b) LC gradient program with 1 minute (upper) and 0.6 minute (bottom) LC column equilibration. (c) Order of LC/MS/MS measurements for evaluation of column equilibration time. (d), (e) Reproducibility of retention time between run 1 and 2, and run 2 and 3. (f) Number of identified peptides in 100 repetitive measurements. (g) Reproducibility of retention time of three identified peptides in 100 repetitive measurements.

**Supplementary Figure S4.**
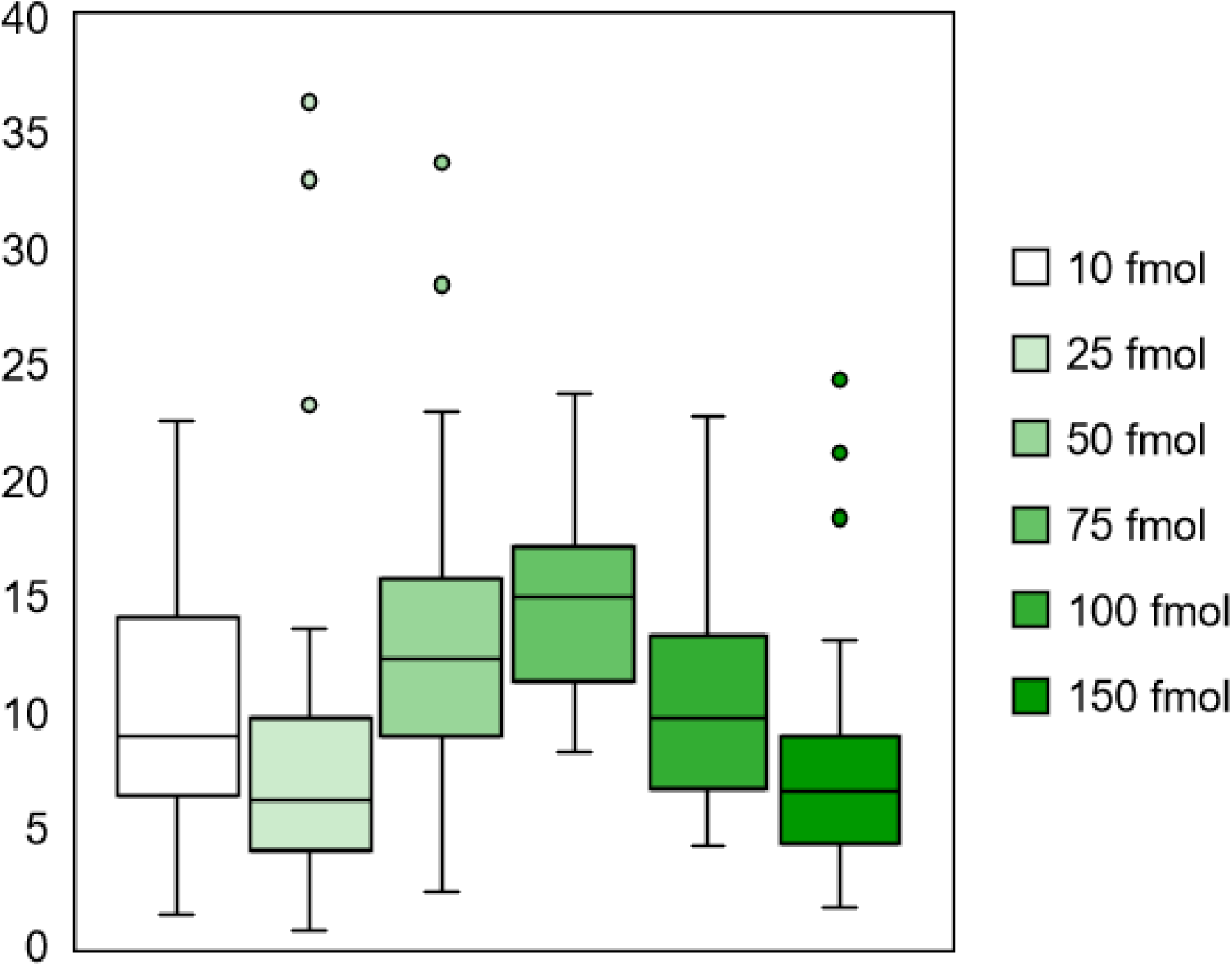
Precision of pY linearity experiments. The CV (%) of pY linearity with matrix experiments.

**Supplementary Table S1.** Experimental methods and results of the experiments of relationship between ESI efficiency and flow rate.

| Flow rate (μL/min) |  | 0.25 | 0.5 | 1 | 5 | 7 | 10 | 50 | 100 | 200 | 500 |
| --- | --- | --- | --- | --- | --- | --- | --- | --- | --- | --- | --- |
| Peptide injection amounts / time (fmol/min) |  | 200 |  |  |  |  |  |  |  |  |  |
| Peptide concentration (fmol/μL) |  | 800 | 400 | 200 | 40 | 28.6 | 20 | 4 | 2 | 1 | 0.5 |
| Relative intensity (%)<br>(/ 250nL/min) | YLYEIAR | 100 | 105.1 | 97.3 | 44.5 | 40.0 | 24.6 | 5.8 | 2.6 | 2.2 | 1.7 |
|  | FALPQYLK | 100 | 73.5 | 61.5 | 35.0 | 29.8 | 24.0 | 7.0 | 3.6 | 2.1 | 1.6 |
|  | YLGYLEQLLR | 100 | 84.6 | 61.3 | 36.9 | 31.3 | 32.7 | 9.1 | 4.3 | 2.5 | 1.5 |
|  | Average | 100 | 87.7 | 73.3 | 38.8 | 33.7 | 27.1 | 7.3 | 3.5 | 2.3 | 1.6 |

**Supplementary Table S2.**
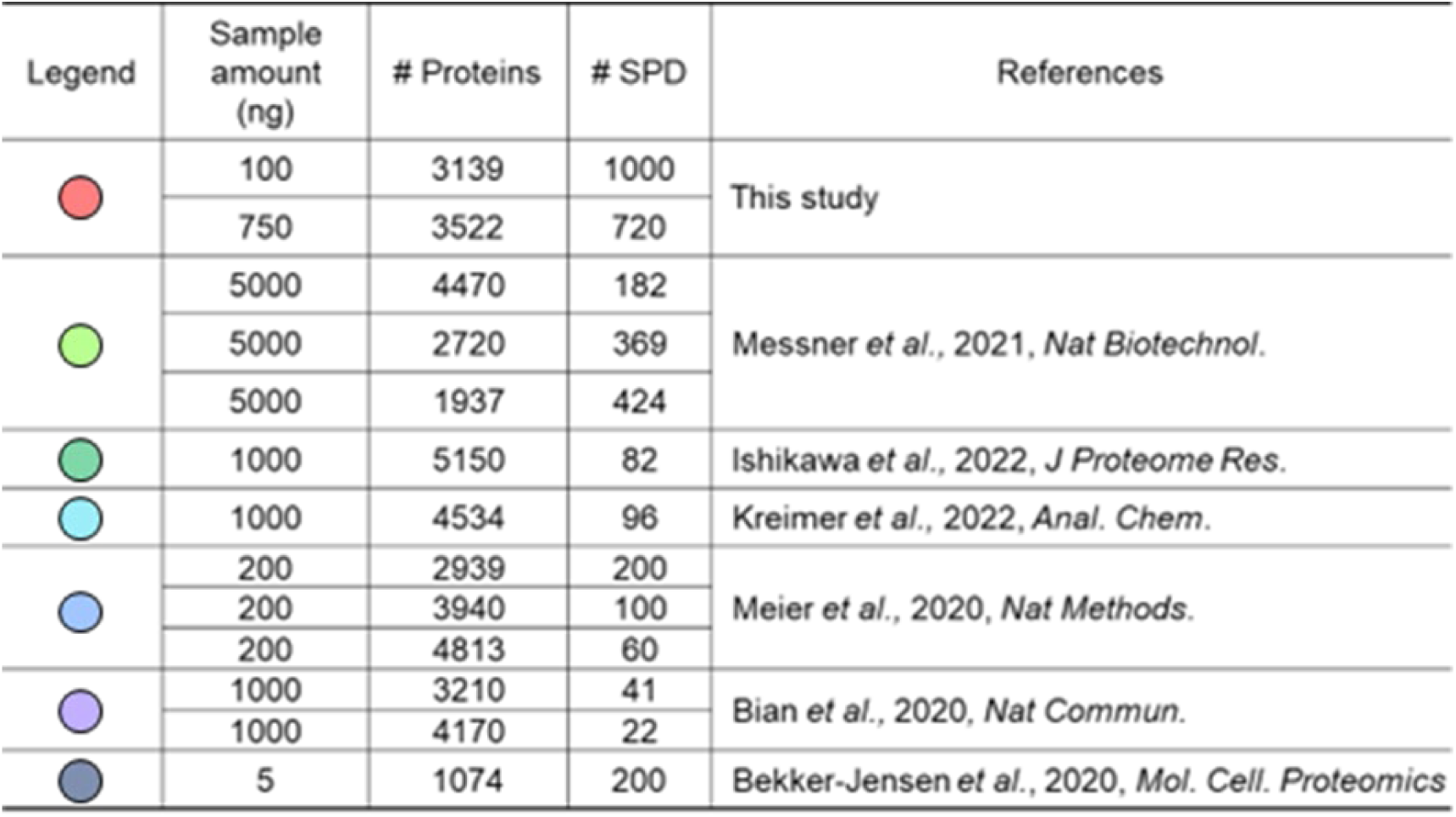
Comparison of our study with other high-throughput proteome analysis methods.

| Legend | Sample amount (ng) | # Proteins | # SPD | References |
| --- | --- | --- | --- | --- |
|  | 100 | 3139 | 1000 | This study |
|  | 750 | 3522 | 720 |  |
|  | 5000 | 4470 | 182 | Messner *et al.*, 2021, *Nat Biotechnol.* |
|  | 5000 | 2720 | 369 |  |
|  | 5000 | 1937 | 424 |  |
|  | 1000 | 5150 | 82 | Ishikawa *et al.*, 2022, *J Proteome Res.* |
|  | 1000 | 4534 | 96 | Kreimer *et al.*, 2022, *Anal. Chem.* |
|  | 200 | 2939 | 200 | Meier *et al.*, 2020, *Nat Methods.* |
|  | 200 | 3940 | 100 |  |
|  | 200 | 4813 | 60 |  |
|  | 1000 | 3210 | 41 | Bian *et al.*, 2020, *Nat Commun.* |
|  | 1000 | 4170 | 22 |  |
|  | 5 | 1074 | 200 | Bekker-Jensen *et al.*, 2020, *Mol. Cell. Proteomics* |

**Supplementary Table S3.** diaPASEF scheme used in this study.

| #MS Type | Cycle Id | Start IM<br>[1/K0] | End IM<br>[1/K0] | Start Mass<br>[m/z] | End Mass<br>[m/z] | CE [eV] |
| --- | --- | --- | --- | --- | --- | --- |
| MS1 | 0 | - | - | - | - | - |
| PASEF | 1 | 1.14 | 1.25 | 976 | 1012 | - |
| PASEF | 1 | 1.03 | 1.14 | 832 | 868 | - |
| PASEF | 1 | 0.92 | 1.03 | 688 | 724 | - |
| PASEF | 1 | 0.81 | 0.92 | 544 | 580 | - |
| PASEF | 1 | 0.7 | 0.81 | 400 | 436 | - |
| PASEF | 2 | 1.17 | 1.28 | 1012 | 1048 | - |
| PASEF | 2 | 1.06 | 1.17 | 868 | 904 | - |
| PASEF | 2 | 0.95 | 1.06 | 724 | 760 | - |
| PASEF | 2 | 0.84 | 0.95 | 580 | 616 | - |
| PASEF | 2 | 0.73 | 0.84 | 436 | 472 | - |
| PASEF | 3 | 1.19 | 1.3 | 1048 | 1084 | - |
| PASEF | 3 | 1.08 | 1.19 | 904 | 940 | - |
| PASEF | 3 | 0.97 | 1.08 | 760 | 796 | - |
| PASEF | 3 | 0.86 | 0.97 | 616 | 652 | - |
| PASEF | 3 | 0.76 | 0.86 | 472 | 508 | - |
| PASEF | 4 | 1.22 | 1.33 | 1084 | 1120 | - |
| PASEF | 4 | 1.11 | 1.22 | 940 | 976 | - |
| PASEF | 4 | 1 | 1.11 | 796 | 832 | - |
| PASEF | 4 | 0.89 | 1 | 652 | 688 | - |
| PASEF | 4 | 0.78 | 0.89 | 508 | 544 | - |

## References

1. Aebersold, R. & Mann, M. Mass-spectrometric exploration of proteome structure and function. Nature 537, 347–355 (2016).

2. Müller, J. B. et al. The proteome landscape of the kingdoms of life. Nature 1–5 (2020).

3. Kawashima, Y. et al. Single-Shot 10K Proteome Approach: Over 10,000 Protein Identifications by Data-Independent Acquisition-Based Single-Shot Proteomics with Ion Mobility Spectrometry. J. Proteome Res. (2022) doi:10.1021/acs.jproteome.2c00023.

4. Muntel, J. et al. Surpassing 10000 identified and quantified proteins in a single run by optimizing current LC-MS instrumentation and data analysis strategy. Molecular Omics 15, 348–360 (2019).

5. Smith, L. M., Kelleher, N. L. & Consortium for Top Down Proteomics. Proteoform: a single term describing protein complexity. Nat. Methods 10, 186–187 (2013).

6. Walsh, C. T., Garneau-Tsodikova, S. & Gatto, G. J., Jr. Protein posttranslational modifications: the chemistry of proteome diversifications. Angew. Chem. Int. Ed Engl. 44, 7342–7372 (2005).

7. Leutert, M., Entwisle, S. W. & Villén, J. Decoding Post-Translational Modification Crosstalk With Proteomics. Mol. Cell. Proteomics 20, 100129 (2021).

8. Bader, J. M., Albrecht, V. & Mann, M. MS-based proteomics of body fluids: The end of the beginning. Mol. Cell. Proteomics 100577 (2023).

9. Zhu, Y. et al. Nanodroplet processing platform for deep and quantitative proteome profiling of 10-100 mammalian cells. Nat. Commun. 9, 882 (2018).

10. Mund, A., Brunner, A.-D. & Mann, M. Unbiased spatial proteomics with single-cell resolution in tissues. Mol. Cell 82, 2335–2349 (2022).

11. Rosenberger, F. A., Thielert, M. & Mann, M. Making single-cell proteomics biologically relevant. Nat. Methods 20, 320–323 (2023).

12. Kelly, R. T. Single-Cell Proteomics: Progress and Prospects. Mol. Cell. Proteomics 22, mcp.R120.002234 (2020).

13. Mund, A. et al. Deep Visual Proteomics defines single-cell identity and heterogeneity. Nat. Biotechnol. 1–10 (2022).

14. Brunner, A.-D., et al. Ultra-high sensitivity mass spectrometry quantifies single- cell proteome changes upon perturbation. (2020).

15. Xiao, Q. et al. High-throughput proteomics and AI for cancer biomarker discovery. Adv. Drug Deliv. Rev. 176, 113844 (2021).

16. Bian, Y. et al. Identification of 7 000–9 000 Proteins from Cell Lines and Tissues by Single-Shot Microflow LC–MS/MS. Anal. Chem. acs.analchem.1c00738 (2021).

17. Bian, Y. et al. Robust, reproducible and quantitative analysis of thousands of proteomes by micro-flow LC-MS/MS. Nat. Commun. 11, 157 (2020).

18. Messner, C. B. et al. Ultra-fast proteomics with Scanning SWATH. Nat. Biotechnol. 1–9 (2021).

19. Szyrwiel, L., Gille, C., Mülleder, M., Demichev, V. & Ralser, M. Speedy-PASEF: Analytical flow rate chromatography and trapped ion mobility for deep high-throughput proteomics. bioRxiv 2023.02.17.528968 (2023) doi:10.1101/2023.02.17.528968.

20. Messner, C. B. et al. Ultra-High-Throughput Clinical Proteomics Reveals Classifiers of COVID-19 Infection. Cell Syst 11, 11–24.e4 (2020).

21. Ishikawa, M. et al. Optimization of Ultrafast Proteomics Using an LC-Quadrupole-Orbitrap Mass Spectrometer with Data-Independent Acquisition. J. Proteome Res. (2022) doi:10.1021/acs.jproteome.2c00121.

22. Kreimer, S. et al. Parallelization with Dual-Trap Single-Column Configuration Maximizes Throughput of Proteomic Analysis. Anal. Chem. 2022.06.02.494601 (2022) doi:10.1021/acs.analchem.2c02609.

23. Bache, N. et al. A Novel LC System Embeds Analytes in Pre-formed Gradients for Rapid, Ultra-robust Proteomics. Mol. Cell. Proteomics 17, 2284–2296 (2018).

24. Bekker-Jensen, D. B. et al. A compact quadrupole-orbitrap mass spectrometer with FAIMS interface improves proteome coverage in short LC gradients. Mol. Cell. Proteomics 19, 716– 729 (2020).

25. Meyer, J. G., Niemi, N. M., Pagliarini, D. J. & Coon, J. J. Quantitative shotgun proteome analysis by direct infusion. Nat. Methods 17, 1222–1228 (2020).

26. Meier, F. et al. Online Parallel Accumulation–Serial Fragmentation (PASEF) with a Novel Trapped Ion Mobility Mass Spectrometer *. Mol. Cell. Proteomics 17, 2534–2545 (2018).

27. Ogata, K. & Ishihama, Y. Extending the Separation Space with Trapped Ion Mobility Spectrometry Improves the Accuracy of Isobaric Tag-Based Quantitation in Proteomic LC/MS/MS. Anal. Chem. 92, 8037–8040 (2020).

28. Meier, F., Park, M. A. & Mann, M. Trapped Ion Mobility Spectrometry (TIMS) and Parallel Accumulation – Serial Fragmentation (PASEF) in Proteomics. Mol. Cell. Proteomics 20, 100138 (2021).

29. Iwasaki, M., Sugiyama, N., Tanaka, N. & Ishihama, Y. Human proteome analysis by using reversed phase monolithic silica capillary columns with enhanced sensitivity. J. Chromatogr. A 1228, 292–297 (2012).

30. Iwasaki, M. et al. One-dimensional capillary liquid chromatographic separation coupled with tandem mass spectrometry unveils the Escherichia coli proteome on a microarray scale. Anal. Chem. 82, 2616–2620 (2010).

31. Wang, H. U., Chen, Z. F. & Anderson, D. J. Molecular distinction and angiogenic interaction between embryonic arteries and veins revealed by ephrin-B2 and its receptor Eph-B4. Cell 93, 741–753 (1998).

32. Borthakur, S., Lee, H., Kim, S., Wang, B.-C. & Buck, M. Binding and function of phosphotyrosines of the Ephrin A2 (EphA2) receptor using synthetic sterile α motif (SAM) domains. J. Biol. Chem. 289, 19694–19703 (2014).

33. Lee, H. & Bennett, A. M. Receptor protein tyrosine phosphatase-receptor tyrosine kinase substrate screen identifies EphA2 as a target for LAR in cell migration. Mol. Cell. Biol. 33, 1430–1441 (2013).

34. Meier, F. et al. diaPASEF: parallel accumulation–serial fragmentation combined with data-independent acquisition. Nat. Methods 17, 1229–1236 (2020).

35. Daigle, T. L. et al. A Suite of Transgenic Driver and Reporter Mouse Lines with Enhanced Brain-Cell-Type Targeting and Functionality. Cell 174, 465–480.e22 (2018).

36. Lein, E. S. et al. Genome-wide atlas of gene expression in the adult mouse brain. Nature 445, 168–176 (2007).

37. Harris, J. A. et al. Hierarchical organization of cortical and thalamic connectivity. Nature 575, 195–202 (2019).

38. Oh, S. W. et al. A mesoscale connectome of the mouse brain. Nature 508, 207–214 (2014).

39. Urisman, A. et al. An Optimized Chromatographic Strategy for Multiplexing In Parallel Reaction Monitoring Mass Spectrometry: Insights from Quantitation of Activated Kinases. Mol. Cell. Proteomics 16, 265–277 (2017).

40. Piehowski, P. D. et al. Automated mass spectrometry imaging of over 2000 proteins from tissue sections at 100-μm spatial resolution. Nat. Commun. 11, 8 (2020).

41. Tsai, C.-F. et al. Surfactant-assisted one-pot sample preparation for label-free single-cell proteomics. Communications Biology 4, 265 (2021).

42. Masuda, T., Tomita, M. & Ishihama, Y. Phase transfer surfactant-aided trypsin digestion for membrane proteome analysis. J. Proteome Res. 7, 731–740 (2008).

43. Rappsilber, J., Mann, M. & Ishihama, Y. Protocol for micro-purification, enrichment, pre-fractionation and storage of peptides for proteomics using StageTips. Nat. Protoc. 2, 1896–1906 (2007).

44. Rappsilber, J., Ishihama, Y. & Mann, M. Stop and go extraction tips for matrix-assisted laser desorption/ionization, nanoelectrospray, and LC/MS sample pretreatment in proteomics. Anal. Chem. 75, 663–670 (2003).

45. Sugiyama, N. et al. Phosphopeptide Enrichment by Aliphatic Hydroxy Acid-modified Metal Oxide Chromatography for Nano-LC-MS/MS in Proteomics Applications*S. Mol. Cell. Proteomics 6, 1103–1109 (2007).

46. Kyono, Y., Sugiyama, N., Imami, K., Tomita, M. & Ishihama, Y. Successive and selective release of phosphorylated peptides captured by hydroxy acid-modified metal oxide chromatography. J. Proteome Res. 7, 4585–4593 (2008).

47. Boersema, P. J., Raijmakers, R., Lemeer, S., Mohammed, S. & Heck, A. J. R. Multiplex peptide stable isotope dimethyl labeling for quantitative proteomics. Nat. Protoc. 4, 484–494 (2009).

